# Restriction of RecG translocation by DNA mispairing

**DOI:** 10.1101/2021.03.31.437879

**Authors:** Zhiqiang Sun, Yaqing Wang, Mohtadin Hashemi, Piero R. Bianco, Yuri L. Lyubchenko

## Abstract

The RecG DNA helicase plays a crucial role in stalled replication fork rescue as the guardian of the bacterial genome. We have recently demonstrated that single-strand DNA binding protein (SSB) promotes binding of RecG to the stalled replication fork by remodeling RecG, enabling the helicase to translocate ahead of the fork. We also hypothesized that mispairing of DNA could limit such translocation of RecG, which plays the role of roadblocks for the fork movement. Here, we used atomic force microscopy (AFM) to directly test this hypothesis and investigate how sensitive RecG translocation is to different types of mispairing. We found that a C-C mismatch at a distance of 30 bp away from the fork position prevents translocation of RecG over this mispairing. A G-bulge placed at the same distance also has a similar roadblock efficiency. However, a C-C mismatch 10 bp away from the fork does not prevent RecG translocation, as 10 bp from fork is within the distance of footprint of RecG on fork DNA. Our findings suggest that retardation of RecG translocation ahead of the replication fork can be a mechanism for the base pairing control for DNA replication machinery.

## Introduction

The interplay between genetic recombination and DNA repair machinery is the key point for genome duplication, which is a highly accurate and processive process (1-3). However, genome duplication can be stopped by DNA template damage, arrested polymerase, and other DNA-protein interactions(4-7). These roadblocks stall the replication fork, and various mechanisms are involved in fork repair and restoring the replication process. RecG is one of the critical proteins that is involved in the restoration process. RecG is a powerful, monomeric, atypical DNA helicase that binds to stalled DNA replication forks and couples DNA unwinding to duplex rewinding, resulting in the extrusion of Holliday junctions(8). RecG binds specifically to the stalled replication fork, rewinds the parental strand DNA utilizing ATP hydrolysis, generates a Holliday junction (HJ), and allows the homologous DNA repair machinery to repair DNA and start the replication process(8-10).

In addition to directly binding to forks and processing them, RecG also physically and functionally interacts with the single-strand binding protein (SSB)(9-15). The physical interaction is utilized by RecG to bind to SSB already positioned on the fork, resulting in the loading of RecG onto duplex DNA, concomitant with helicase remodeling(16). After RecG’s binding mode to the DNA fork being remodeled, RecG is able to spontaneously translocate in front of the fork DNA over distances as large as 200bp (17). We were able to directly visualize such thermally driven and ATP-independent translocation of RecG with time-lapse high-speed AFM (HS-AFM). These findings suggest that mispairing of the DNA duplex in front of the fork can limit the RecG translocation distance, thereby sending a signal to the replication machinery about potential problems with the DNA duplex ahead of the fork.

To investigate this hypothesis, we created forks with duplex imperfections in the parental dsDNA arm. The duplex imperfections include a C-C mismatch or a G-bulge, positioned either 10 or 30 bp ahead of the fork (Fig. 1). While RecG can bind directly to forks, we have previously shown that SSB enhances the interaction of the helicase with forks. Therefore, each fork used in this study has a gap in the leading strand arm (the preferred fork for RecG), but this arm is 69nt in length to ensure efficient SSB binding, as before (10,16-18). The binding of RecG to each fork in the absence and presence of SSB was separately assessed using AFM. The interaction with SSB initiates RecG remodeling, defined by the elevated affinity of RecG to the fork. Insertions of the G-bulge and C-C mismatch 30 bp away from the fork resulted in a decrease of the translocation distance from 48 ± 13 bp to 26 ± 11 bp and 26 ± 10 bp (S.D.), respectively. The C-C mismatch placed 10 bp from the fork has less of an effect on the translocation distance, allowing RecG to translocate over distances 30 ± 11 bp rather than stall at 10 bp. The explanation for this trend is given by the simulated structure of RecG-DNA complex(19), which showed that RecG covers 23 bp on the DNA duplex region. Mispairing at the 30 bp distance hinders the protein translocation, while the mismatch 10 bp away from the fork does not. These results reveal a novel function of RecG, in which it selects the DNA duplex quality of parental arm of the fork.

**Figure 1.**
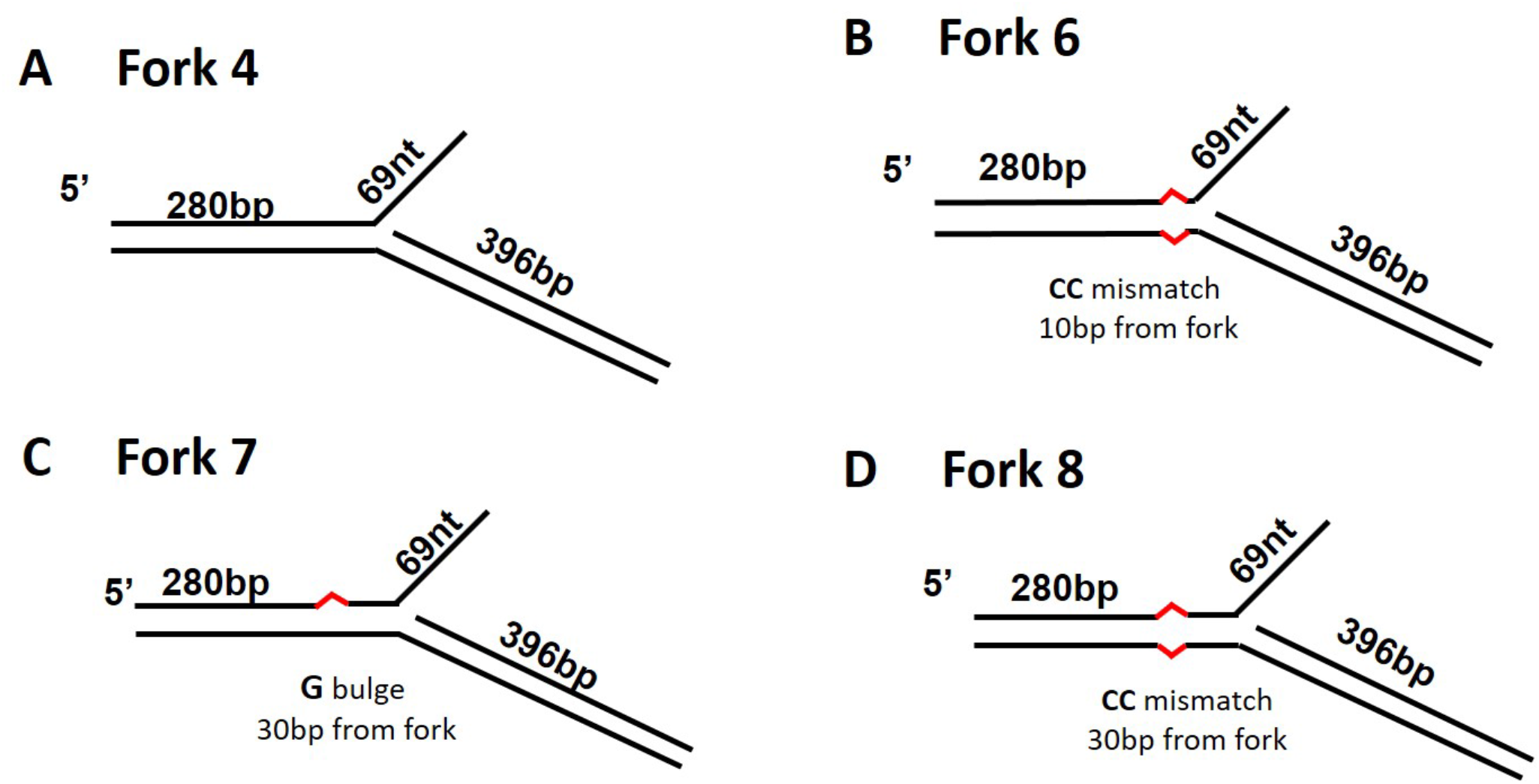
The scheme of fork substrates. (A) The F4 construct contains 69nt ssDNA between the two perfectly paired DNA duplexes (280bp and 396bp). (B, C, D) The F6, F7 and F8 constructs, respectively, containing mismatches or bulges at different positions of the parental strand. (B) F6 contains a C-C mismatch 10 bp away from the fork. (C) F7 contains a single G bulge 30 bp upstream of the fork. (D) F8 contains a C-C mismatch 30 bp away from the fork.

## Results and discussion

### Fork DNA constructs

Our constructs that mimic stalled DNA replication forks, with and without additional damage, are schematically shown in Fig. 1. Construct F4 represents a stalled replication fork with a gap in the nascent leading strand flanked by asymmetric DNA duplex regions (280 and 396 bp in length). Previously, we have used F4 to study RecG binding, which allowed us to distinguish between parental and lagging strand arms of the fork in AFM images(16). This type of replication fork is similar to the substrate in the crystallographic studies of *T. maritima* RecG-DNA complexes^(19)^. Constructs F6, F7, and F8 are similar to F4, except they contain a mismatch or bulge in the parental duplex region. F6 contains a C-C mismatch at 10 bp ahead of the fork position. F7 contains a single G-bulge on the parental arm positioned 30 bp upstream of the fork, while F8 contains a C-C mismatch at the same position. The C-C mismatch can change the local structure of DNA, including global helical bending, opening, and minor groove width in the center of the helix(20,21). The dynamics (breathing frequency) of local DNA of C-C mismatch suggests that C-C mismatch is stronger than other kind of mismatches(22). The G bulge in the F7 construct contains an extra G on one ssDNA, and there is two C on each sides of G, which makes it less stable than others(23). The mismatch or bulge positions were designed to be within the thermal sliding distance of SSB bound to a fork (16,17) (24).

### SSB facilitates RecG binding to the fork constructs

Previously, we found that SSB facilitates RecG binding to fork substrates(16). Here, we tested whether SSB can facilitate the binding of RecG to different fork constructs containing mispaired regions. The experiments for all substrates were performed in parallel. As shown in Fig. 2, when SSB and RecG are added sequentially to the reaction mixture, both proteins can be observed bound to the same DNA (indicated by the arrows). As reported previously(16), SSB and RecG can be readily discerned; this is displayed in the insets of Fig. 2, with the larger feature being SSB (blue arrow) and the smaller one being RecG (red arrow). Furthermore, for most DNA molecules with both proteins bound, SSB and RecG are at distinct positions, and seldom colocalize. There are some DNA molecules with only a single feature, and analysis shows that these are SSB (Fig. S1).

**Figure 2.**
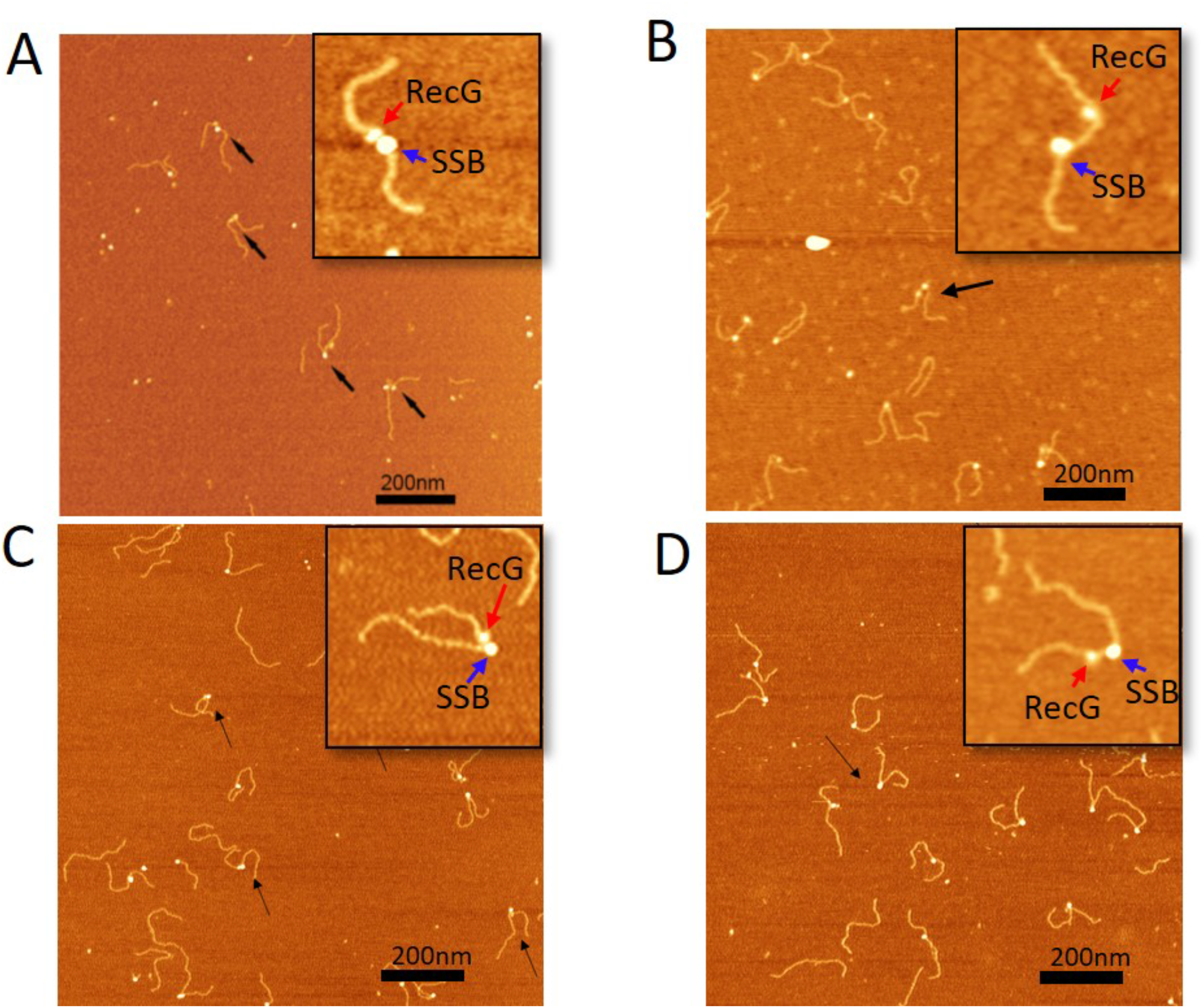
AFM images of SSB and RecG bound to various fork substrates are shown in (A) for F4, (B) for F6, (C) for F7 and (D) for F8. Double-feature complexes are indicated by arrows. The scale bar is 200 nm. The insets (200 nm x 200 nm) are the zoomed images of typical double-feature complexes; the blue arrows point to SSB, and the red arrows point to RecG protein.

To assess the ability of SSB to load RecG onto the designed forks, we measured the yield of protein-DNA complexes in RecG only and SSB-RecG experiments. Results show that the yield of RecG complexes is higher when SSB is added first compared to RecG only (Fig. 3). The number of DNA-protein complexes varies from 50 to 80. These data also show that when the duplex imperfection is positioned 10 bp from the fork (Fork 6), binding of RecG is inhibited by 2-fold, regardless of the presence of SSB. In contrast, when the duplex imperfection is positioned 30 bp away from the fork, RecG binding or SSB loading of the helicase is unaffected, with F7 and F8 substrates producing data similar to F4. This suggests that when the imperfection is within the RecG binding footprint with the fork, RecG binding is inhibited. Additionally, this also suggests that the initial loading site of RecG by SSB encompasses 10 base pair from the fork.

**Figure 3.**
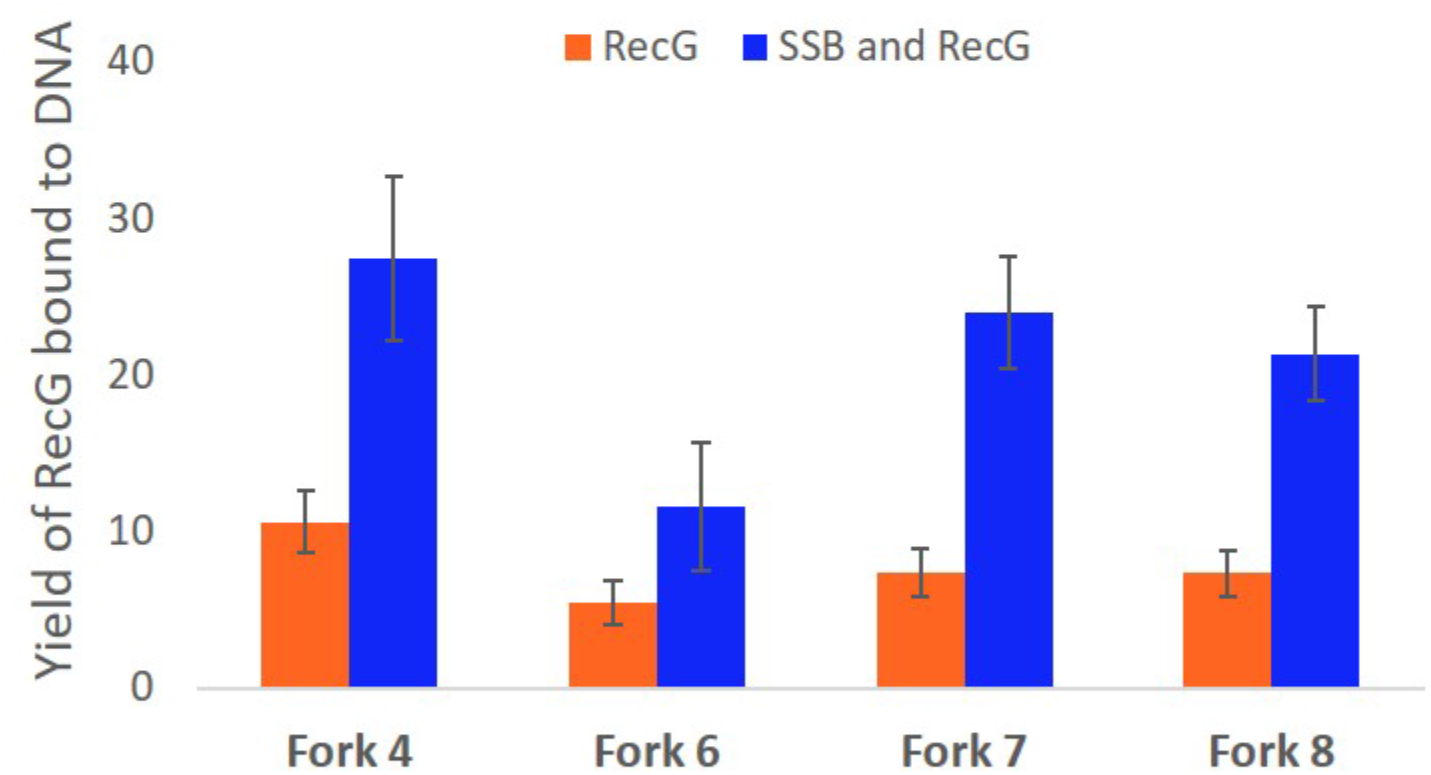
Effect of SSB on the binding of RecG to various forks. Yields of SSB-RecG complexes with replication fork substrates were calculated in the absence (orange bars) and the presence of SSB (blue bars). In the absence of SSB, the binding efficiency of RecG on F4, F6, F7 and F8 substrates are 10.6±1.9%, 5.4±1.2%, 7.4±1. 5% and 7.3±1.5%, respectively. SSB increases the RecG binding efficiency to the fork DNA substrates to 27.4±5.3%, 11.6±4.1%, 24±3.6% and 21.3±3%, respectively.

### The mispairing limits the translocation of RecG

SSB can enhance the binding efficiency of RecG on all four different fork substrates; therefore, we investigated whether SSB can remodel RecG on all fork substrates and allow RecG to translocate over the arms of the fork. To do this, we mapped positions of SSB and RecG on F4, F6, F7, and F8 substrates, and the data is shown in Fig. 4 (A-D), respectively. SSB and RecG positions were measured relative to the same DNA end, usually the end of the parental strand (short arm). Since SSB binds specifically to the ssDNA, the position of SSB was set to 0. When RecG translocates to the parental strand, the position of RecG has a negative value (below SSB in the graph). When RecG translocates to the lagging strand, the position of RecG has a positive value (above SSB in the graph). Similar to our previous findings(16), when RecG binds to the fork DNA in the presence of SSB, RecG appears on the duplex arms of the fork. Furthermore, when translocating, RecG prefers the parental arm on all fork substrates. In control experiments in the absence of SSB, RecG binds precisely to the fork position (Fig. S2).

**Figure 4.**
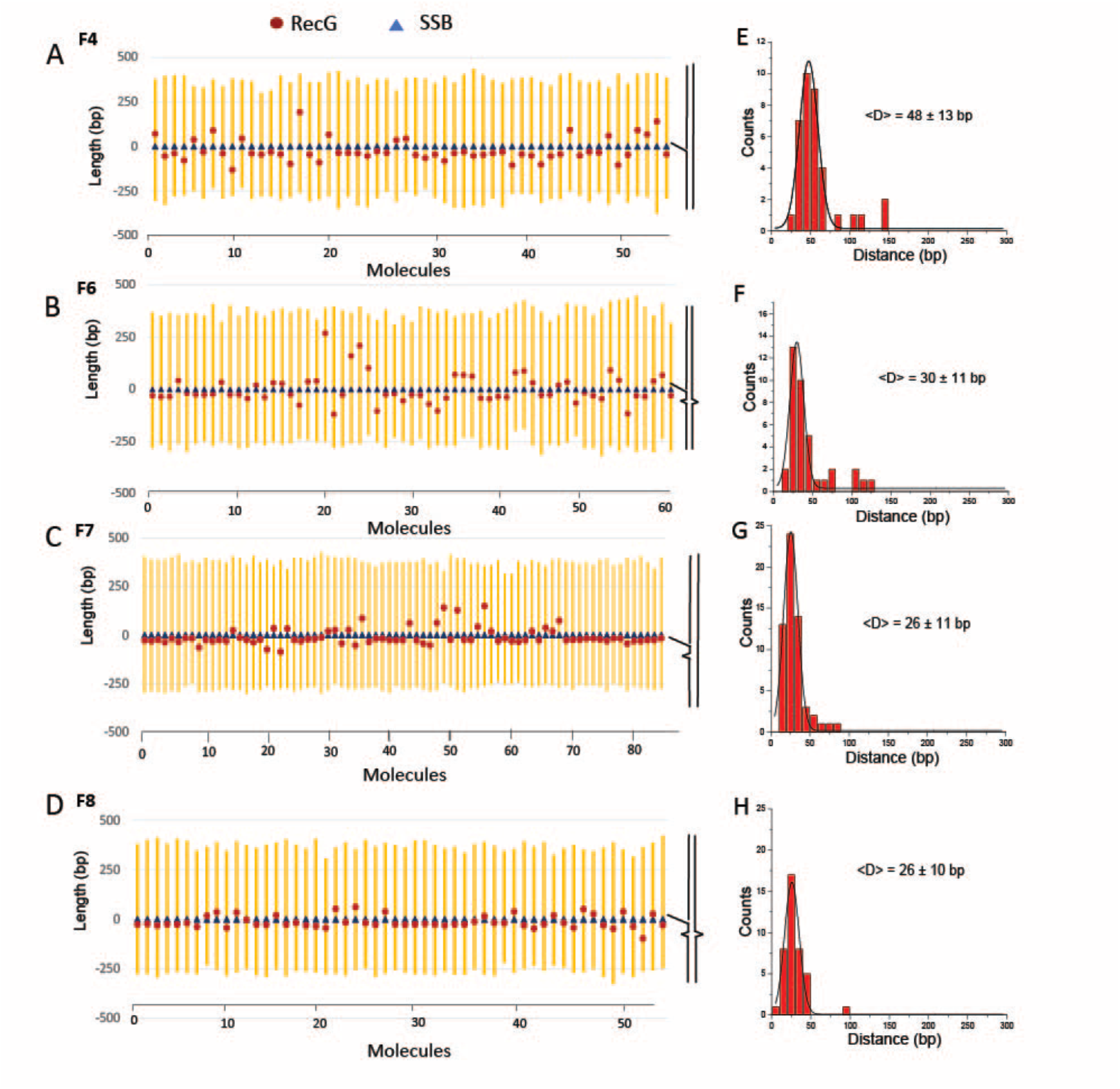
Mapping of RecG positions relative to the SSB binding site. The map is constructed relative to the position of SSB bound to ssDNA of the replication fork. On the maps of SSB is indicated with triangles and RecG with circles. Frames A, B, C and D show positions of RecG on the DNA substrates F4, F6, F7 and F8, respectively. In 75% of cases, RecG binds to the shorter duplex segment (parental) of the F4 construct. While in F6, F7 and F8 construct RecG binds to the longer segment (parental) of the DNA duplex 60%, 75% and 74% of the time. Histograms (E-H) show distributions of distances between RecG (on parental arms) and SSB measured which correspond to respective maps. Histograms were approximated with Gaussians and the values corresponding to maxima on these distributions are inserted in the histograms.

Translocation distance on the parental arm was investigated due to the fact that all mismatches are located on the parental arm of F6, F7, and F8. These translocation values are shown as histograms of RecG positions in Fig. 4 (E-H) for DNA substrates F4, F6, F7, and F8, respectively. The data show that RecG translocation distance is shorter on the fork substrates with C-C mismatch or G-bulge. For construct F4, RecG is loaded onto the parental strand arm in 75% of the cases, with the translocation distance being 48 ± 13 bp. The arm preference for F6, F7, and F8 constructs are 60%, 75%, and 74% for parental arm, respectively, while translocation distances are 30 ± 11 bp, 26 ± 11 bp and 26 ± 10 bp. The shorter translocation distance, compared to on F4, suggests that the C-C mismatch and G-bulge can limit the translocation of RecG on the parental arm. Additionally, the translocation distances of RecG on F7 and F8 are very similar and match the position of the bulge and mismatch location (30 bp from fork position). The translocation distance of RecG on F6 also decreased compared to that on F4, but all the RecG were found at positions beyond 10 bp away from the fork.

We hypothesized that RecG has a large footprint on the fork DNA, so the mismatch on F6 will be within the binding regions and cannot work as a roadblock. Fig. 5 shows that the footprint of RecG on the duplex DNA (green + red) is 23 bp, which is obtained from our previous RecG structure and simulations with a fork DNA substrate(19). This suggests that when the position of the DNA mismatch is more than 23 bp away from the fork, it should not affect the binding of RecG to the fork. The mismatch at 10 bp away from the fork on F6 is within the footprint of RecG. Thus, it can decrease the binding efficiency of RecG to the fork but cannot limit the translocation of RecG when RecG is bound to the fork. Indeed, the yield of RecG on F6, even in the presence of SSB, is lower than that on the other substrates. Interestingly, the translocation distance of RecG on the parental arm of F6 is still less than that on F4. This suggests that the 10 bp mismatch location also affects the translocation of RecG, but to a lesser extent and through a possibly different mechanism compared to the mismatch located 30 bp from the fork position.

**Figure 5.**
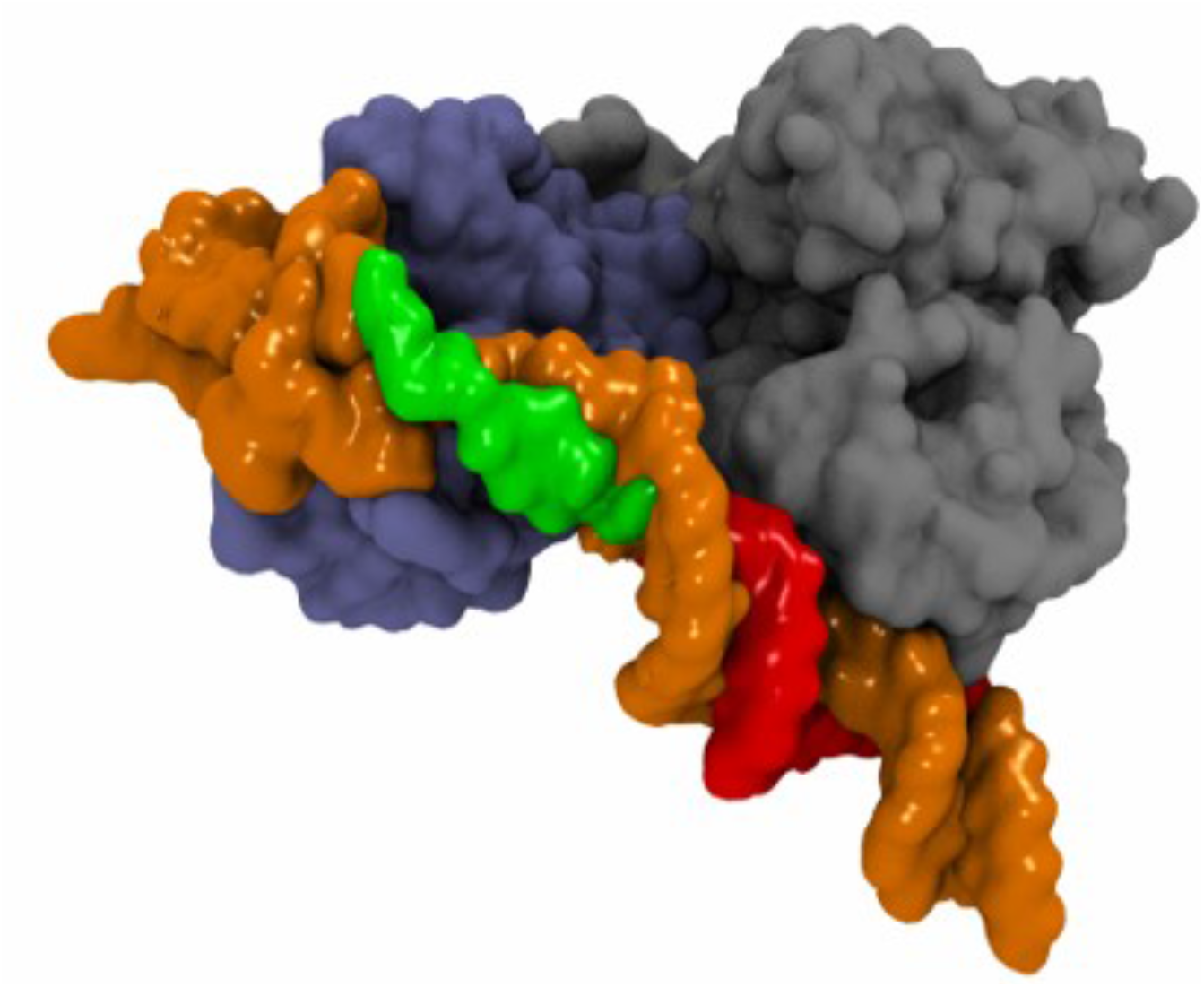
Snapshot of the RecG interacting with a stalled replication fork substrate after a 150 ns MD simulation. RecG wedge domain is colored purple, and the helicase domain is colored gray. DNA is colored orange and the segment between the wedge domain and helicase domain is in green, and the DNA interacting with the helicase domain is in red. The size of the green DNA is 10 nt while the red is 13 nt.

Compared with narrow distributions of translocation distances on parental arms, the distances between SSB and the RecG on the lagging arm vary in a wide range (Fig. S3). There is no preferred translocation distance of RecG on the lagging strand. This suggests that while the translocation of RecG on the parental strand is limited by the mismatches 30 bp from the fork, the lagging strand does not impose such a limit on the translocation of RecG since there is no mismatches on lagging strand arm. While most RecG translocated short distances from the fork, the data in Fig. S4 show that RecG can pass over the mispairings and translocate to distances up to ∼300 bp on both arms of fork DNA, which was also observed in our previous findings(17). On F6 substrate, with the C-C mismatch 10 bp away from the fork, the frequency of long-range translocation is greater than that on the other fork substrates. This may be due to the mismatch located 10 bp from the fork, which does not block the translocation, allowing RecG to move far beyond the mismatch position. On F7, majority of the RecG appeared on the lagging strand (Fig. S3 and S4) and far away from SSB (more than 50 bp). On F8, there were only a few RecG molecules translocating more than 50 bp.

Overall, our data demonstrate that mispairings can act as roadblocks for spontaneous RecG translocation. Due to a large footprint of RecG on the DNA replication fork, the mispairing can be detected by RecG at distances beyond 20 bp. In those cases, the presence of RecG in front of the replication fork may act as a signal for the replication machinery to stop and for the repair process to start. The location of RecG in the proximity of the replication fork facilitates its participation in the fork regression process, which is the major function of RecG.

## Methods

### Protein preparations

RecG protein was purified as described previously (25). Briefly, the protein was eluted using a linear salt gradient (10–1000 mM NaCl), with RecG eluting between 250 and 360 mM NaCl, on a 100 ml Q-Sepharose column equilibrated in buffer A [20 mM Tris-HCl (pH 8.5), 1 mM EDTA, 1 mM DTT, 10 mM NaCl]. The pooled fractions were then subjected to heparin FF and hydroxylapatite chromatography, as previously described (26). Pooled fractions from the hydroxylapatite column were dialyzed overnight into S buffer [10 mM KPO_4_ (pH 6.8), 1 mM DTT, 1 mM EDTA and 100 mM KCl]. The protein was applied to a 1 ml MonoS column and eluted using a linear KCl gradient (100–700 mM) with RecG eluting at 350 mM KCl. The fractions containing RecG were pooled and dialyzed overnight against storage buffer [20 mM Tris-HCl (pH 7.5), 1 mM EDTA, 1 mM DTT, 100 mM NaCl and 50% (v/v) glycerol]. The protein concentration was spectrophotometrically determined using an extinction coefficient of 49,500 M^−s^ cm^−c(26)^.

Single-strand DNA-binding protein was purified from strain K12ΔH1Δtrp as previously described (27). The concentration of purified SSB protein was determined at 280 nm using ε = 30,000 M^-1^ cm^-1^. The site size of SSB protein was determined to be 10 nucleotides per monomer by monitoring the quenching of the intrinsic fluorescence of SSB that occurs on binding to ssDNA, as described previously (15,28).

### Preparation of fork DNA substrates

The fork DNA substrates were assembled from two duplexes and the core fork segment, similar to our previous methodology(16). Briefly, the core fork was made by annealing for short oligoes. Then the cork fork was ligated with two long duplexes on both sides. The core fork segment varied between the substrates to include one of the two the mismatches. The core segment of F4 substrate was assembled from O30 (TCATCTGCGTATTGGGCGCTCTTCCGCTTCCTATCT), O31 (TCGTTCGGCTGCGGCGAGCGGTATCAGCTCACTCATA), O32 (GCTTATGAGTGAGCTGATA CCGCTCGCCGCAGCCGAACGACCTTGCGCAGCGAGTCAGTGAGATAGGAAGCGGAA GAGCGCCCAATACGCAGA), and O33 (CACTGACTCGCTGCGCAAGGTCGTTCGGCTGCGGCGAGCGGCTA ACATCTGGGTTTTCATTCTTTGGGTTTCACTTTCTCCAC). To assemble the core segment of F6, O33 was replaced by O35 (CACTGACTCCCTGCGCAAGGCTAACAGCATCACACACATTAACAATTCTAACATCTGGGTTTTCATTCTTTGGGTTTCACTTTCTCCAC). To assemble the core segment of F7, O30 was replaced by O30-bulge (TCATCTGCGTATTGGGCGCTCTTCGCGCTTCCTATCT). For F8, O30 was replaced by O30-mismatch (TCATCTGCGTATTGGGCGCTCTTCCCCTTCCTATCT). All other segments for F6, F7 and F8 are the same as for F4 substrate. All oligonucleotides were obtained from IDT (Integrated DNA Technologies, Inc. Coralville, Iowa, USA).

### Preparation of DNA-protein complexes

*SSB-RecG-DNA complexes*. DNA was mixed with the SSB tetramer in a molar ratio of 1:2 (DNA:SSB) and incubated in binding buffer [10mM Tris-HCl (pH 7.5), 50 mM NaCl, 5 mM MgCl_2_, 1 mM DTT] for 10 min at room temperature. RecG (4:1 molar ratio to DNA) was added into the mixture and incubated for additional 30 min. The final molar ratio of DNA:SSB:RecG was 1:2:4, and the final DNA concentration was 2 nM.

### The sample preparation and AFM imaging

APS functionalized mica was used as the AFM substrate for all experiments. Briefly, freshly cleaved mica was incubated in a 167 µM aqueous solution of 1-(3-aminopropyl)silatrane (APS) for 30 min and rinsed thoroughly with deionized water as described in(16). Five microliters of the sample were deposited onto the APS mica for 2 min, rinsed with deionized water, and dried with a gentle argon gas flow. Images were acquired using tapping mode in air on a MultiMode 8, Nanoscope V system (Bruker, Santa Barbara, CA) using TESPA probes (320 kHz nominal frequency and a 42 N/m spring constant) from the same vendor.

### Data analysis

The AFM images were analyzed using the FemtoScan Online software package (Advanced Technologies Center, Moscow, Russia), which enables precise tracing of the DNA molecules. The position of each protein relative to the end of the short arm on the DNA substrate was measured. The total DNA length was then measured continuously to the other end of the DNA substrate. The yield of complexes was calculated from the ratio of protein-DNA complexes to the total number of DNA molecules.

## Funding

The work was supported by the National Institutes of Health (R01 GM100156 to YLL and PRB, R01 GM096039, R01GM118006 to YLL).

## Author contributions

YLL, ZS, YW and MH conceived, designed the experiments. ZS purified the DNA constructs and collected AFM images, PRB provided all proteins, MH performed computational modeling. All authors wrote the manuscript.

## Acknowledgement

Thank Thomas Stormberg for the discussion and writing of the manuscript.

## Completing financial interests

The authors declare no competing financial interests.

## Supporting Information

Figures S1-S4.

